# Intracellular pH controls Wnt signaling downstream of glycolysis in the vertebrate embryo

**DOI:** 10.1101/481259

**Authors:** Masayuki Oginuma, Yukiko Harima, Fengzhu Xiong, Olivier Pourquié

**Affiliations:** Department of Genetics, Harvard Medical School and Department of Pathology, Brigham and Women’s Hospital, Boston, MA, USA; Harvard Stem Cell Institute, Harvard University, Cambridge, MA USA

**Author notes:** Correspondence to: O.Pourquié. These authors contributed equally to this work (Masayuki Oginuma, Yukiko Harima).

## Abstract

Formation of the body of vertebrate embryos proceeds sequentially by posterior addition of tissues. While this process depends on aerobic glycolysis acting upstream of Wnt signaling in tail bud cells, the molecular details of this regulation are unknown. Here we used chicken embryos and human tail bud-like cells differentiated *in vitro* from iPS cells to show that glycolysis acts by increasing the intracellular pH (pHi) of tail bud cells. This promotes β-catenin acetylation leading to Wnt signaling activation and the choice of a mesodermal fate at the expense of the neural fate in tail bud precursors. Our data suggest that by increasing the pHi of tail bud cells, aerobic glycolysis creates a favorable chemical environment for non-enzymatic acetylation of β-catenin, ultimately triggering Wnt signaling.

## Main text

Cells of the tail bud and posterior Presomitic Mesoderm (PSM) exhibit a high level of aerobic glycolysis which is reminiscent of the metabolic status of cancer cells experiencing Warburg effect (*1*). We previously reported that 2-day chicken embryos exhibit a posterior to anterior gradient of extracellular pH (pHe), with the lowest pH found in the tail bud where glycolysis is most active (*2*). Treatment of such embryos with the glycolysis inhibitor 2 Deoxy-D-glucose (2DG) or culture in glucose-free medium results in a uniform higher pHe consistent with a role for glycolysis in the acidification of the extracellular environment at the posterior end of the embryo (*2*). In normal differentiated adult cells, the intracellular pH (pHi) is usually around 7.2 whereas the pHe is generally around 7.4. In most cancer cells, the regulation of pHe and pHi is significantly different from normal cells (*3*): whereas pHe is lower (6.7-7.1) than in normal adult cells due to lactic acid excretion, pHi is higher (>7.4) compared to normal cells (*3*). To examine whether PSM and tail bud cells also exhibit an inverted pHe-pHi gradient, we measured the pHi in these tissues in chicken embryos. We used the ratiometric pH sensor pHluorin, a modified pH-sensitive Green Fluorescent Protein (GFP) which has been previously used to analyze pHi *in vivo* (*4, 5*). To test whether pHluorin can accurately report on pHi differences in the chicken embryo, we electroporated a pHluorin construct in the anterior primitive streak region (which contains the PSM precursors) of stage 4HH chicken embryos (*6*). One day after electroporation, embryos were incubated in buffers of different pH (pH5.5, pH6.5, pH7.5) with the protonophores Nigericin and Valinomycin. This allows the cells’ pHi to equilibrate to the buffer pH (*5*). We next measured the average emission signal ratio at 488/405 nm in electroporated cells located in the tail bud and the PSM using confocal imaging and single cell 3D segmentation. As expected, we observed that the ratio decreased as the buffer pH increased (fig S1), indicating that the pHluorin reporter electroporated *in vivo* can report on intra-cytoplasmic pH changes in embryos. We next measured the 488/405 nm ratio in electroporated embryos in the absence of the protonophores. We observed a posterior to anterior gradient of pHluorin 488/405 nm signal ratio in the majority of wild type embryos (n=11/13) (Fig. 1A-B). With some local variability, the tail-bud cells show a higher fraction of 405 nm excitable pHluorin whereas anterior PSM cells show a higher fraction of 488 nm excitable pHluorin (Fig. 1A-B). Thus, anterior PSM cells have higher intra-cytoplasmic acidity (lower pHi) compared to posterior cells.

**Figure 1:**
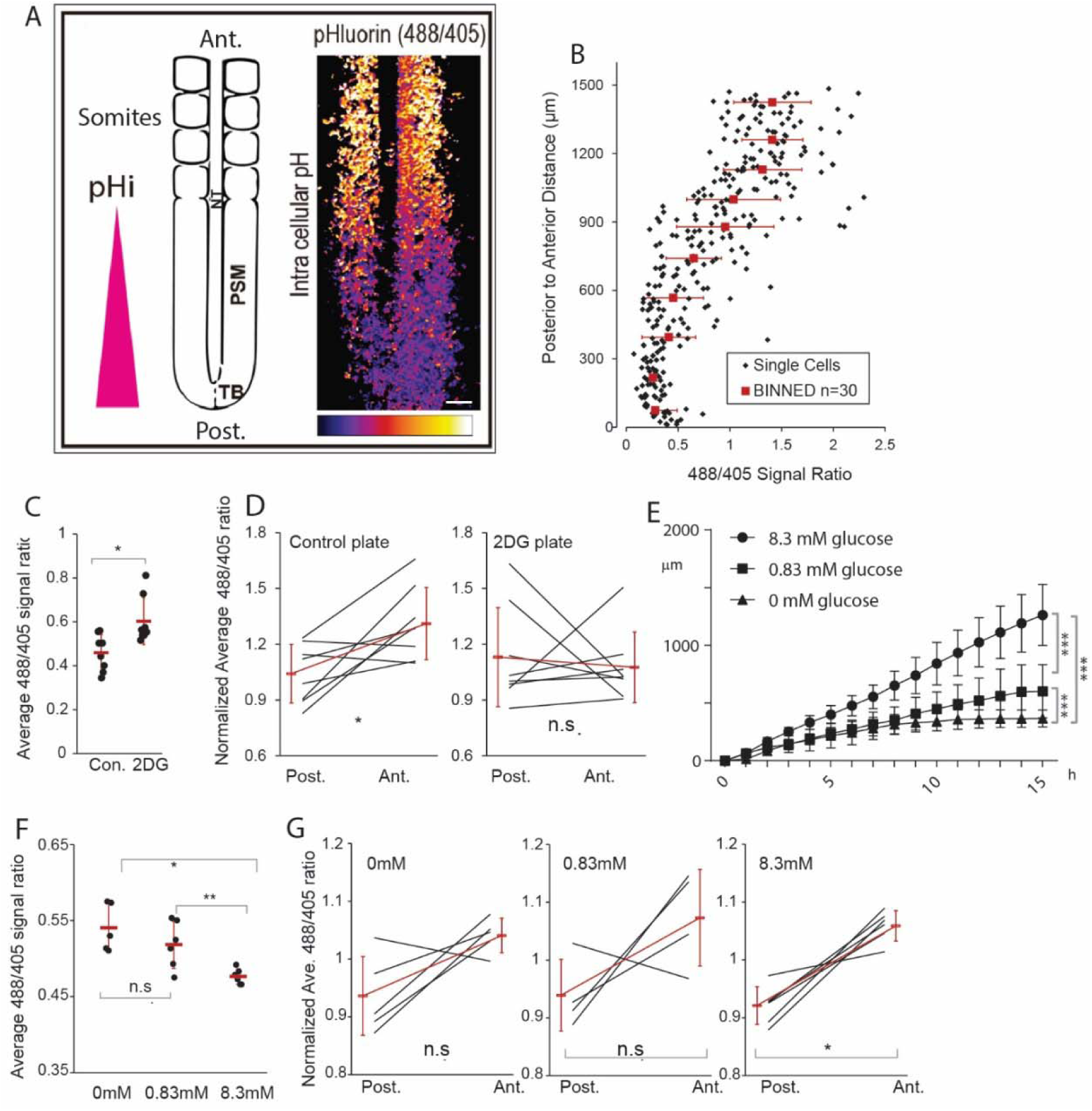
An antero-posterior intra-cytoplasmic pH gradient in the tail bud and posterior PSM is regulated by glycolysis. (A) Intra-cytoplasmic pH (pHi) in the posterior part of a 2-day old chicken embryo expressing pHluorin in the Tail Bud (TB) and Presomitic Mesoderm (PSM). Fluorescence intensity is shown by pseudocolor image (Fire color) using image J (right panel). Yellow signal indicates more acidic regions. Ventral view. NT: neural tube. Ant.: anterior, Post.: posterior. Scale bar, 100 μm. (B) 488/405 nm ratio in individual cells in a representative embryo (n=3). Average values are calculated every 30 segmented cells along the AP axis. Error bars are S.D. (C) Average 488/405 nm ratio of the PSM region in control (Con) (n=8) and 2DG (n=8) embryos in EC culture.*p=0.01, statistical significance assessed by t-test. (D) Average 488/405 nm ratio in anterior and posterior PSM in control (n=8) and 2DG (n=8) embryos. Each black line represents data from one embryo. Normalized 488/405 nm ratios of 40 posterior-most and 40 anterior-most cells are averaged to generate the ends of each line. Red lines are average (±SD) of different embryos. Post.: Posterior. Ant.: Anterior. * p=0.02; n.s. p=0.69 (paired t-tests). Scale bar: 100 µm. (E) Increase in axis length (elongation) measured over time using time lapse microscopy (mean ±SD) in the chemically defined medium at pH7.2 with 8.3mM glucose (circles, n=6), 0.83mM glucose (squares, n=3) and 0 mM glucose (triangles, n=6). Normal development is observed in the chemically defined medium although axis elongation is slower compared to normal Embryo Culture medium (EC) (85 μm/h vs 170 μm/h in normal EC culture). Statistical significance was assessed with two way Anova followed by Tukey’s multiple comparison test. ***p<0.001. (F) Average 488/405 nm ratio of the PSM region of 2-day chicken embryos cultured in chemically defined medium at pH7.2 with 8.3 mM glucose (n=6), 0.83 mM glucose (n=5) and 0 mM glucose (n=6). In each sample, the average ratio (black dot) is obtained by averaging all cells (n varies between 50-250 per embryo). The samples were pooled for an average ratio +/- SD (red bars) and subjected to a t-test. *p=0.01; **p=0.001; n.s. p>0.05. (G) Normalized ratios of 40 cells on the posterior and the anterior ends of the PSM were averaged (ends of each black line). Embryos were cultured in chemically defined medium at pH7.2 with 8.3 mM glucose (n=6), 0.83 mM glucose (n=5) and 0 mM glucose (n=6). The samples were pooled for an average ratio +/- SD (ends of the red line) and subjected to a paired t-test. *p=0.002

We next explored whether glycolysis plays a role in the control of pHi in PSM cells. Treating embryos electroporated with pHluorin in the PSM with 2DG *ex ovo* increased the overall 488/405 nm ratio of the tail bud and posterior PSM (Fig. 1C), suggesting a global increase of intracellular acidity. Importantly, electroporated embryos grown on 2DG plates show no anterior-posterior gradient of 488/405 nm ratios compared to the embryos grown on control plates (Fig. 1D). We observed that culturing 2-day old chicken embryos in agar plates containing only PBS and 8.3 mM glucose at pH7.2 is sufficient to sustain normal development (*2, 7*) (Fig. 1E). Decreasing the glucose concentration to 0.83 mM and 0 mM in the minimal medium resulted in a dose-dependent slowing down of axis elongation similar to that seen with 2DG treatment (*2*) (Fig. 1E, fig. S2). We next cultured embryos in which pHluorin was electroporated in the PSM precursors in these low glucose conditions and measured the 488/405 nm signal ratio in the PSM. Decreasing the glucose concentration led to a dose-dependent increase of the 488/405 nm ratio, suggesting a decrease in pHi as observed when inhibiting glycolysis with 2DG (Fig. 1F). No significant pHi gradient could be evidenced in embryos cultured in low glucose conditions (Fig. 1G). Thus, decreasing glycolytic activity in cultured embryos leads to an increase of intracytoplasmic acidity in PSM cells and abolishes the anterior-posterior gradient of pHi. These data therefore support the idea that the observed spatial pH differences come from differential glycolytic activity along the PSM.

We next investigated the role of pHi in the control of PSM cell differentiation *in vivo*. Embryos electroporated with pHluorin were cultured *ex ovo* in minimal medium buffered at different pH with NaHCO3 for 3 hours and we measured the 488/405 nm signal ratio in the PSM (Fig. 2A). The signal ratio in the posterior PSM decreased when the pH of the medium increased indicating that exposing embryos to a lower pHe leads to a corresponding acidification of the pHi in the PSM (Fig. 2B). When embryos were cultured at pH6.0, axis elongation was significantly slowed down (Fig. 2C, fig. S2). Culturing the embryos at pH5.3 led to a stronger phenotype with axis elongation rapidly stalling (Fig. 2C, fig. S2). These phenotypes are similar to those observed when inhibiting glycolysis with 2DG (*2*) or with low glucose concentrations (Fig. 1E). Remarkably, embryos exposed to a low pH buffer switched back to control medium after elongation arrest resumed normal development (fig. S2). Thus, the developmental arrest observed after lowering the pH is not due to irreversible damage to the embryo. Together, these experiments suggest that appropriate regulation of pHi is essential for normal elongation of the embryonic axis.

**Figure 2:**
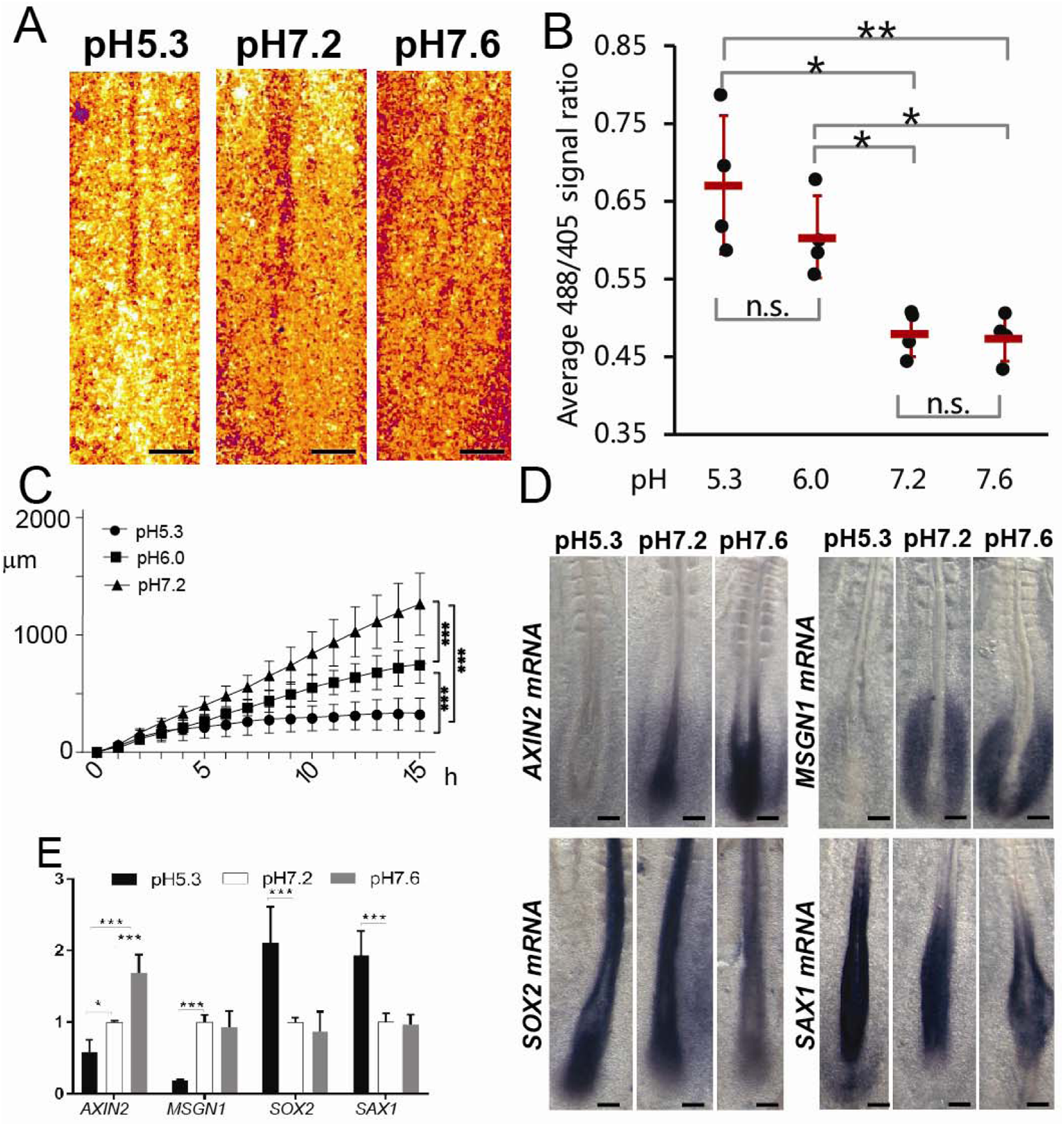
pHi regulates Wnt activity in the posterior region of chicken embryos. (A) Fluorescence intensity at 488 nm showing the posterior part of a 2-day old chicken embryo electroporated at stage 4HH with pHluorin and cultured in chemically defined medium with 8.3 mM glucose at different pH. Fluorescence intensity is shown by pseudocolor image (Fire color) using image J. Yellow signal indicates acidic regions. Ventral view, anterior to the top. Scale bar, 100 μm. (B) Average 488/405 nm ratio of 2-day chicken embryos cultured in chemically defined medium with 8.3 mM glucose at different pH (pH5.3: n=4, pH6.0: n=4, pH7.2: n=4, pH7.6: n=4). In each sample, the average ratio (black dot) is obtained by averaging all cells (n varies between 50-250 per embryo). The samples were pooled for an average ratio +/- SD (red bars) and subjected to a t-test. *p<0.01; **p<0.001; n.s. p>0.05. (C) Increase in axis length (elongation) measured over time using time lapse microscopy (mean ±SD) in 2-day chicken embryos cultured in chemically-defined medium with 8.3 mM glucose at different pH: pH7.2 (triangles, n=6), pH6.0 (squares, n=5) and pH5.3 (circles, n=8). Statistical significance was assessed with two way Anova followed by Tukey’s multiple comparison test. ***p<0.001. (D) Whole-mount *in situ* hybridization of 2-day chicken embryos cultured for 10 h in chemically defined medium at different pH and hybridized with *AXIN2* (pH5.3: n=6, pH7.2: n=6, pH7.6: n=4), *MSGN1* (pH5.3: n=4, pH7.2: n=3, pH7.6: n=3), *SOX2* (pH5.3: n=6, pH7.2: n=4, pH7.6: n=5), and *SAX1* (pH5.3: n=6, pH7.2: n=4, pH7.6: n=5). Ventral view, anterior to the top. Scale bar, 100 μm. (E) qPCR analysis of *MSGN1, SAX1*, *SOX2* and *AXIN2* expression in the posterior region of 2-day chicken embryos cultured for 10 h in chemically defined medium with 8.3 mM glucose at different pH (n=4 for each gene). Error bars represent SD. Statistical significance was assessed with unpaired two-tailed student t-test. * p<0.05, ***p<0.001.

In the tail bud, glycolysis is required for Wnt activation, which is necessary for axis elongation and for paraxial mesoderm production from the bi-potential neuro-mesodermal precursor (NMP) population (*2, 8, 9*). Blocking glycolysis with 2DG *in vivo* phenocopies Wnt signaling loss of function (*2*). This results in most NMPs to differentiate to a neural SOX2/SAX1-positive fate, ultimately leading to elongation arrest (*2*). Our data indicate that glycolysis is necessary to maintain a high pHi in the PSM/tail bud region and to support axis elongation, thus raising the possibility that glycolysis regulates Wnt signaling via the control of pHi. To test this hypothesis, we examined how altering tail bud pHi impacts the expression of the canonical Wnt target *AXIN2* using *in situ* hybridization and qPCR (Fig. 2D-E). Strikingly, culturing embryos in a low pH buffer decreases *AXIN2* expression while culture at high pH results in stronger expression (Fig. 2D-E). A similar behavior is observed for *MSGN1,* another Wnt/β-catenin target specifically expressed in the posterior PSM (Fig. 2D-E) (*10*). In contrast, the neural genes *SOX2* or *SAX1* showed an opposite behavior, being upregulated at low pH (Fig. 2D-E). Together, these results suggest that higher pHi favors canonical Wnt signaling, promoting the paraxial mesoderm fate from NMPs, while lower pHi inhibits Wnt, promoting the neural fate. Thus, these data argue that glycolysis could play a role in controlling Wnt signaling in tail bud cells by maintaining a high pHi.

We recently reported a protocol to differentiate human pluripotent cells to the paraxial mesoderm fate *in vitro* (*11, 12*). Treatment of human iPS cells with the GSK3β inhibitor CHIR99021 (Chir) in combination with the BMP inhibitor LDN193189 (LDN) (CL medium), induces the majority of cells to differentiate to a SOX2-BRACHYURY positive NMP-like fate after day 1 (*13*). After day 2, more than 95% of the cells differentiate to a MSGN1-positive posterior PSM fate (*11-13*). In the following days, cells continue to differentiate, expressing the anterior PSM/dermomyotome marker PAX3 around day 6, and they can be subsequently differentiated to the myogenic lineage (*11, 12*). We first examined whether the human iPS cells differentiating to the paraxial mesoderm lineage recapitulate the metabolic status of the tail bud and PSM cell populations observed *in vivo* in chicken and mouse embryos (*2, 14*). We detected a decrease of lactate dehydrogenase (*LDHB*) and of lactate production after day 2 in culture (Fig. 3A). This suggests that differentiating PSM cells downregulate glycolysis as described in the developing mouse and chicken embryos (*2, 14*). In the mouse PSM, nuclear β-catenin is only found in the posterior region which exhibits high glycolytic activity (*15*). In the differentiating iPS cultures examined by immunohistochemistry, nuclear β-catenin localization (which is generally associated to Wnt activation (*16*)) peaked at day 2 (Fig. 3B-C) (*13*). This correlated with the expression of the PSM Wnt targets *AXIN2, TBX6* and *MSGN1* measured by qPCR, which peaked around day 2-3 of differentiation (Fig. 3D-F) (*17-19*). Thus, PSM cells exhibit an increased Wnt response during the early phase of PSM maturation *in vitro*, as observed in the posterior PSM *in vivo (20)*. We next examined the pHi of iPS cells differentiating to the PSM fate *in vitro* using the pH sensitive ratiometric dye BCECF (*5*). We observed a higher pHi (>7.5) at day 2 progressively decreasing in sync with Wnt signaling after day 3 (Fig. 3G). Thus, the decrease of glycolytic and Wnt activity parallels a progressive decrease of pHi in human PSM cells differentiating *in vitro*. These data show that the metabolic status of mouse and chicken PSM cells is largely recapitulated in human iPS-derived PSM cells differentiated *in vitro*.

**Figure 3:**
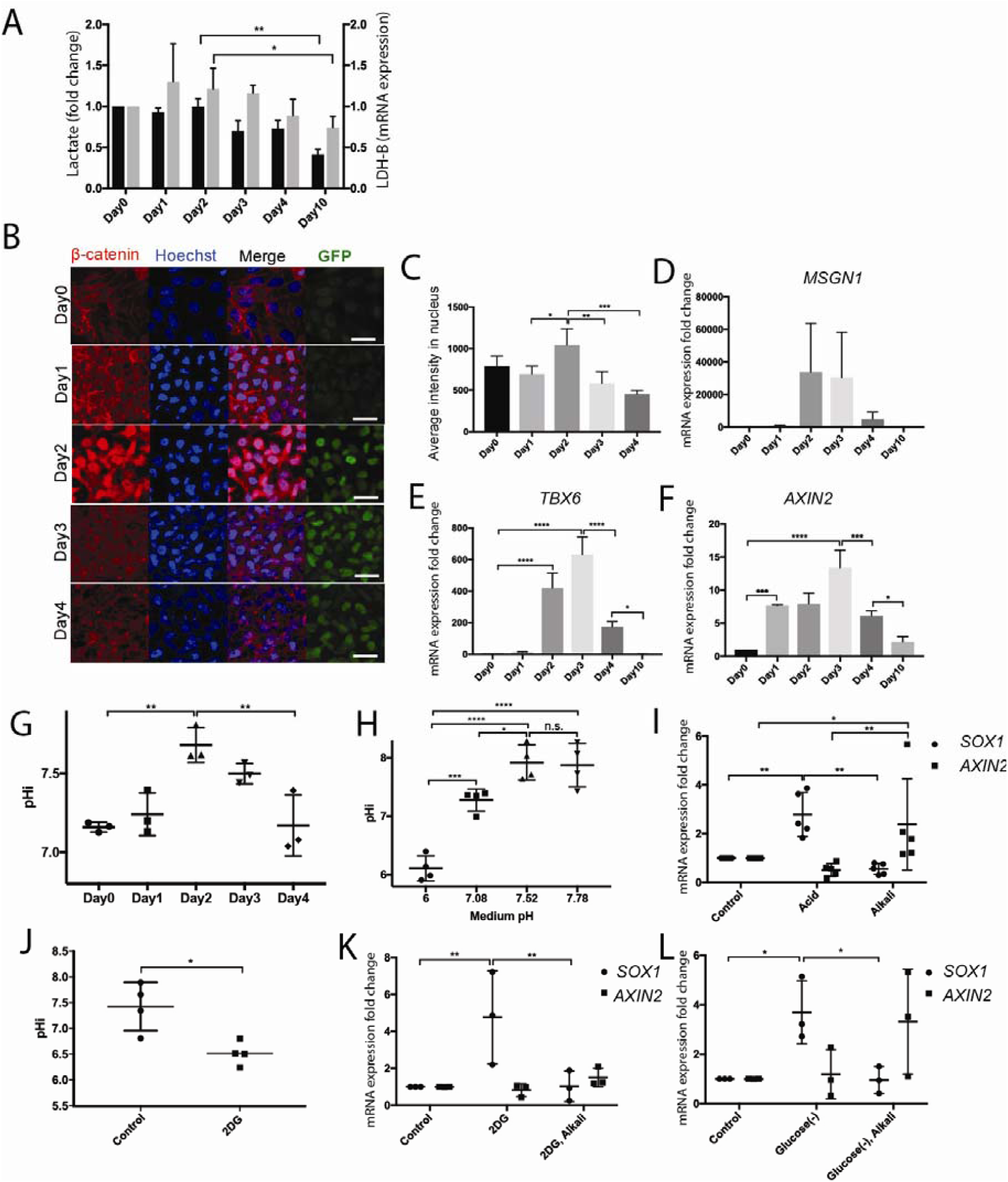
Decrease of pHi and glycolysis levels during *in vitro* differentiation of human iPS- derived PSM cells. (A) Dynamics of lactate production (black bar, fold change of lactate production normalized to total protein content) and *LDH-B* mRNA expression level (gray bar, normalized to *actin* mRNA levels) in human iPS cells differentiated to a PSM fate *in vitro* in CL medium. Graphs show triplicate experiments. Values were normalized by the results of differentiation at day0. Error bars represent SD (n=3). Statistical significance was assessed with two-way ANOVA followed by Tukey’s multiple comparisons test: *p < 0.05; **p< 0.01. (B) Immunohistochemistry showing the dynamic expression of β-catenin and Venus (YFP) proteins in human MSGN1-Venus iPS reporter cells differentiated to the PSM fate *in vitro* in CL medium (n=3). Hoechst labeling of the nuclei is shown in blue. Scale bar: 30 μm (C) Quantification of the intensity of nuclear localization of β-catenin shown in (B) using Fiji. Error bar represent SD (minimum n= 3 experiments per condition). Statistical significance was assessed with one-way ANOVA followed by Tukey’s multiple comparisons test: *p < 0.05; **p< 0.01; ***p< 0.001. (D-F) qPCR analysis comparing the expression level of *MSGN1* (D), *TBX6* (E) and *AXIN2* (F) of human iPS cells differentiating to the PSM fate *in vitro* in CL medium. Values were normalized by the results of differentiation at day0. Error bars represent SD (n=3). Statistical significance was assessed with one-way ANOVA followed by Tukey’s multiple comparisons test: *p < 0.05; **p< 0.01; ***p< 0.001; ****p< 0.0001. (G) Analysis of the pHi using the BCECF dye in human iPS cells differentiating to the PSM fate *in vitro* in CL medium. Error bars represent SD (n=3). Statistical significance was assessed with one-way ANOVA followed by Tukey’s multiple comparisons test. **p < 0.01. (H) Analysis of the pHi using the BCECF dye in day 2 human iPS cells differentiated to the PSM fate *in vitro* in CL medium and cultured for 3 h in CL culture medium at different pH (n=4 for each condition). Error bars represent SD. Statistical significance was assessed with two-way ANOVA followed by Tukey’s multiple comparisons test: *p < 0.05; ***p< 0.001; ****p<0.0001; n.s. p> 0.5 (I) Comparison of *SOX1* and *AXIN2* mRNA expression in day 2 human iPS cells differentiated to a PSM fate *in vitro* in CL medium and cultured for 3 h in CL medium at different pH (Control: pH7.0-7.2), Acid: pH6.3-6.5; Alkali: pH7.5-7.8) (n>=5 for each condition). Error bars represent SD. Statistical significance was assessed with two-way ANOVA followed by Tukey’s multiple comparisons test: *p < 0.05; **p< 0.01. (J) Analysis of the pHi using the BCECF dye in human iPS cells cultured for one day in CL medium followed by 24 h in CL medium with 2DG (n=4 for each condition). Error bars represent SD (n=4). Statistical significance was assessed with unpaired two-tailed test: *p < 0.05. (K-L) Comparison of *SOX1* and *AXIN2* mRNA expression in human iPS cells cultured for one day in CL medium followed by 24 h in CL culture medium with 2DG (K) or in glucose free CL medium (L) at normal pH or in alkaline conditions (pH7.5-7.8) (n=3 experiments for each condition). Error bars represent SD. Statistical significance was assessed with two-way ANOVA followed by Tukey’s multiple comparisons test: *p < 0.05; **p< 0.01.

We observed that culturing iPS cells in CL medium at different pH can predictably change the pHi of cells (Fig. 3H). By qPCR analysis, we showed that decreasing the pHi by culturing cells in acidic conditions at a stage equivalent to the NMP stage (day 1) increased the expression of the neural marker *SOX1* and decreased the expression of *AXIN2* as was observed *in vivo* (Fig. 3I). Compared to acidic conditions, cells exposed to basic conditions (pH7.5 to 8) showed significantly decreased expression of *SOX1* and increased expression of *AXIN2* (Fig. 3I). Therefore, these data suggest that higher pHi promotes the paraxial mesoderm fate from tail bud-like cells differentiating *in vitro* whereas lower pHi favors the neural fate in these cells.

We also noted that reducing glycolytic activity by treating differentiating iPS cells *in vitro* with 2DG decreases the pHi of cells (Fig. 3J). This is accompanied by an increase in *SOX1* expression (Fig. 3K). This effect could be significantly rescued by incubating cells cultured in 2DG-containing medium at a higher pH (Fig. 3K). In these conditions, *SOX1* expression was decreased while *AXIN2* was increased (Fig. 3K). A similar rescue was observed when cells were cultured in glucose free-medium (Fig. 3L). Thus, our *in vivo* and *in vitro* data suggest that pHi acts as a second messenger regulating Wnt signaling downstream of glycolysis in tail bud-like cells.

The dynamic regulation of Wnt signaling in differentiating iPS cells is somewhat unexpected as it occurs despite the constant presence of the Wnt activator Chir in the culture medium. Chir inhibits the kinase GSK3-β which phosphorylates β-catenin and targets it for degradation (*16, 21*). This results in β-catenin stabilization allowing it to bind members of the Lef/TCF family which are required for shuttling β-catenin to the nucleus and for transcriptional activation of the Wnt targets (*16*). In whole extracts from cultures of human iPS-derived PSM cells, we observed a stable expression of β-catenin and its non-phosphorylated (active) form between day 1 to day 4, consistent with the expected effect of GSK3-β inhibition by Chir (Fig. 4A). Thus, the down-regulation of nuclear β-catenin and Wnt targets observed in spite of Chir-treatment suggests that events downstream of β-catenin stabilization in the cytoplasm are dynamically regulated in PSM cells differentiating *in vitro*. One such candidate is K49 β-catenin acetylation, which acts as a switch controlling mesodermal vs neural gene activation in mouse embryonic stem cells (*22*). To test whether K49 β-catenin acetylation is associated with the peak of Wnt activation *in vitro*, we used an antibody which specifically detects K49 acetyl β-catenin. We observed a dynamic expression of K49 acetyl β-catenin peaking at day 1-3 *in vitro* (Fig. 4A). We analyzed the cytoplasmic and nuclear fractions of differentiating iPS cultures by western blot and observed a peak of nuclear K49 acetyl β-catenin around day 2 coincident with the peak of Wnt activation (Fig. 4B). Switching differentiating iPS cells to acidic conditions for 3 hours at day 2 led to a decrease of K49 acetylated β-catenin whereas exposure to alkaline conditions increased acetylated β-catenin levels compared to control (Fig. 4C). In human iPS cells differentiating to PSM, K49 acetylated β-catenin expression decreased in the absence of glucose (Fig. 4D). In all these different conditions, levels of non-phosphorylated and total β-catenin remained stable (Fig. 4C-D). Stimulating global acetylation by treating differentiating cells with acetate, led to an increase of K49 β-catenin acetylation while global levels of β-catenin and its non-phosphorylated form remained stable (Fig. 4E). In addition, acetate treatment led to an upregulation of *MSGN1* and a decrease in *SOX1* expression, consistent with the PSM fate choice triggered by Wnt signaling activation (Fig. 4F). Thus, our data suggest that K49 β-catenin acetylation plays a role in the regulation of Wnt signaling during PSM differentiation *in vitro*.

**Figure 4:**
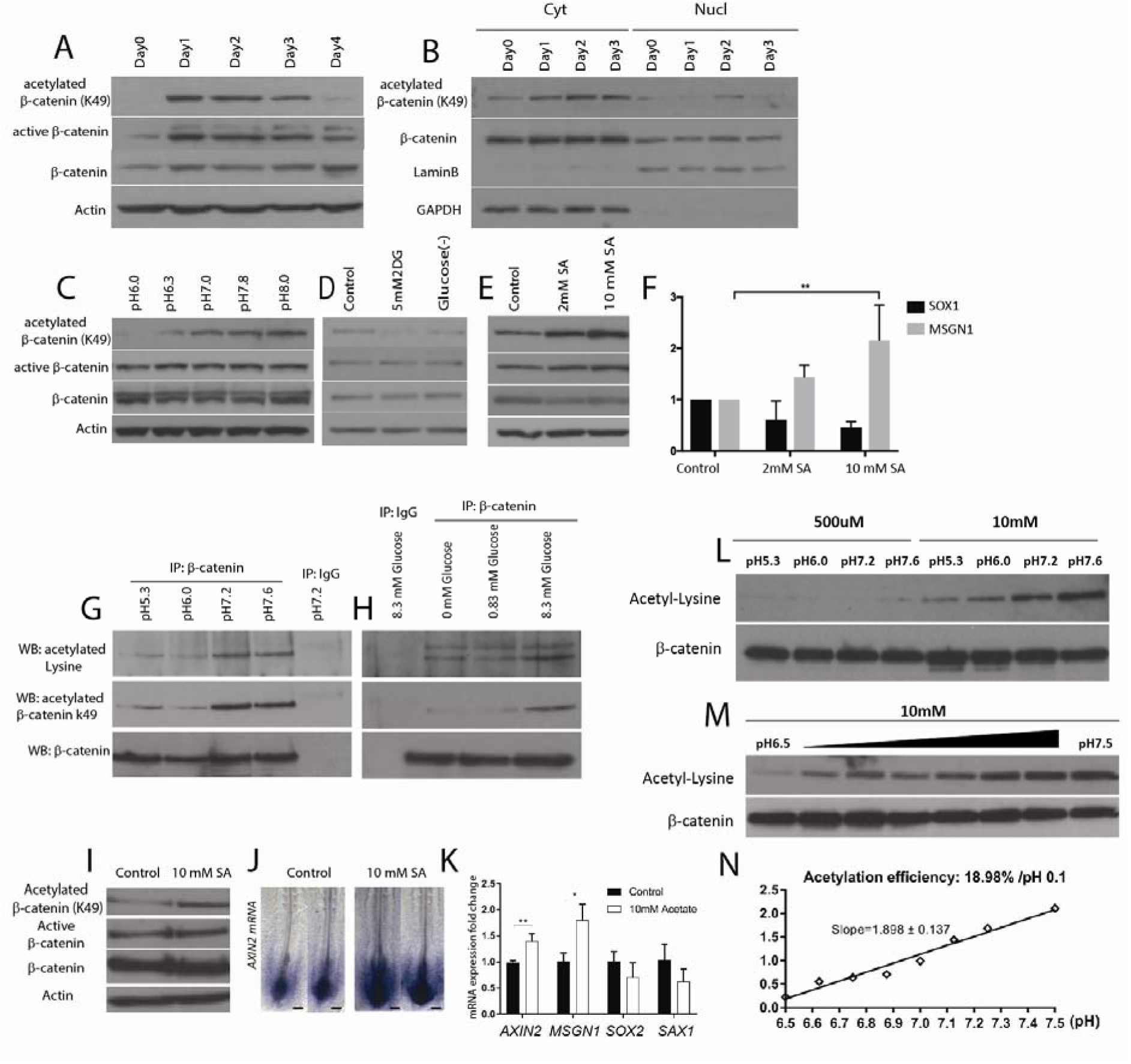
Regulation of β**-catenin acetylation by glycolysis and pHi**. (A) Western blot analysis of whole cell extracts prepared from hiPS cells differentiated to PSM *in vitro* in CL medium, using anti-acetylated K49 β-catenin, anti-active β-catenin, anti-actin and anti-β-catenin antibodies (n=3). (B) Western blot analysis using anti-acetylated K49 β-catenin, anti-laminB, anti-GAPDH and anti-β-catenin antibodies of cytoplasmic (Cyt) and nuclear (Nucl) extracts prepared from hiPS cells differentiating to PSM *in vitro* in CL medium (n=3). (C-E) Western blot analysis using anti-acetylated K49 β-catenin, anti-active β-catenin, anti-actin and anti-β-catenin antibodies. (C) Whole cell extracts of day 2 hiPS cells differentiated to PSM *in vitro* in CL medium and cultured at different pH for 3h (n=4). (D) Whole cell extracts of day 2 hiPS cells differentiated to PSM in CL medium *in vitro* and cultured for 6 h in 2DG and glucose-free CL medium (n=3). (E) Whole cell extracts of day 2 hiPS cells differentiated to PSM *in vitro* in CL medium and treated with Sodium Acetate (SA) for 24 h (n=3). (F) qPCR analysis of *SOX1* and *MSGN1* mRNA expression in day 2 hiPS cells differentiated to PSM *in vitro* and treated with Sodium Acetate (SA) in CL medium for 24 hours. Error bars represent SD (n=3). Statistical significance was assessed with two-way ANOVA followed by Tukey’s multiple comparisons test: **p< 0.01. (G) Immunoprecipitations with an anti β-catenin antibody of extracts of 2-day chicken embryos cultured in chemically defined medium at different pH. Western blot analysis using anti-cetylated lysine, anti-acetylated K49 β-catenin and anti-β-catenin antibodies. IgG: control immunoprecipitation. (n=3). (H) Immunoprecipitations with an anti β-catenin antibody of extracts of 2-day chicken embryos cultured in chemically-defined medium at different glucose concentrations. Western blot analysis using anti-acetylated lysine, anti-acetylated K49 β-catenin and anti-β-catenin antibodies. IgG: control immunoprecipitation. (n=4). (I) Western blot analysis using anti-acetylated K49 β-catenin, anti-active β-catenin, anti-actin and anti-β-catenin antibodies of whole cell extracts of 2-day chicken embryos cultured in chemically defined medium with 0 or 10mM Sodium Acetate (SA) for 10 h. (n=3). (J) Whole-mount *in situ* hybridization of 2-day chicken embryos cultured for 10 h in chemically defined medium with 0 or 10 mM Sodium Acetate (SA) and hybridized with *AXIN2* (control n=5, 10 mM Sodium Acetate n=7). Scale bar, 100 μm. (K) qPCR analysis of *AXIN2, MSGN1, SOX2* and *SAX1* expression in the posterior region of 2- day chicken embryos cultured for 10 h in chemically defined medium with 0 or 10 mM Sodium Acetate. Statistical significance was assessed with unpaired two-tailed student t-test. *p<0.01, **p<0.001. Error bars represent SD. (n=4). (L) Western blot analysis using anti-acetylated lysine antibody (top) and anti-β-catenin antibodies (bottom). Recombinant β-catenin protein was incubated with acetyl-CoA sodium salt in PBS at different pH conditions for 3 h at 37C. (n=3). (M) Western blot analysis using anti-acetylated lysine antibody (top) and anti-β-catenin antibodies (bottom). Recombinant β-catenin protein was incubated with acetyl-CoA sodium salt in PBS at pH6.5∼pH7.5 for 3 h at 37C. (n=3). (N) Quantification of acetylated lysine and β-catenin intensity in (M) using Fiji. Acetylation rate is calculated from the slope of graph.

Using an anti-acetyl-lysine antibody, we found that β-catenin is also acetylated in the PSM of chicken embryos (Fig. 4G). Exposure of 2-day chicken embryos to low pH (pH5.3 or 6.0) led to decreased acetylated β-catenin levels in the PSM while exposure to high pH (pH7.6) does not alter acetylated β-catenin levels compared to control embryos (Fig. 4G). Similar variations in acetylation levels were also observed with the antibody specific for K49 acetylation of β-catenin (Fig. 4G). In contrast, levels of active non-phosphorylated and of total β-catenin were stable when embryos were exposed to buffers of different pH (Fig. 4G, fig. S3). This indicates that acetylation of β-catenin is specifically affected by pH changes as observed *in vitro*. Reduction of acetylated and K49 acetyl but not total or non-phosphorylated β-catenin levels was also observed when reducing glucose concentration in embryos cultured in minimal medium (Fig. 4H). Conversely, treatment of 2-day chicken embryos with Sodium Acetate led to an upregulation of K49 β-catenin acetylation but not of total or non-phosphorylated β-catenin levels (Fig. 4I). This also resulted in an upregulation of *AXIN2* and *MSGN1* (Fig. 4J-K). Together these data show that β-catenin acetylation levels in the chicken embryo PSM can be modulated by pHi, glucose or acetate concentration independently of total and non-phosphorylated β-catenin levels. Together, our results suggest that in the tail bud, the regulation of pHi downstream of glycolysis can tune β-catenin acetylation to control Wnt activation.

How can upregulation of the pHi lead to an increase of β-catenin acetylation? The pHi can control the protonation of specific histidines in proteins acting as pH sensors, leading to changes in the protein properties (*3*). In some epithelial cells, the pHi level can regulate β-catenin stability by controlling its binding to β-TrCP and hence its degradation (*23*). While acetylation of protein substrates such as histones is largely regulated by HATs and HDACs enzymes, non-enzymatic acetylation of proteins has also been demonstrated in many different cell types (*24*). This chemical addition of acetyl-residues to cellular proteins can be regulated by the pHi of cells (*25*). This raises the interesting possibility that the high pHi observed in PSM cells downstream of glycolysis might provide favorable chemical conditions for non-enzymatic β-catenin acetylation. To test this possibility, we performed an *in vitro* reaction, incubating recombinant β-catenin protein with acetyl-CoA sodium salt in PBS at pH5.0, pH6.3, pH7.2, and pH7.6 for 3 hours at 37C. Using an anti-acetyl lysine antibody, a clear dose-dependent increase of β-catenin acetylation was detected when the pH was raised (Fig. 4L). Strikingly, even small pH increments within a physiological range (pH6.5 to pH 7.5) led to significant dose-dependent increase of β- catenin acetylation (Fig. 4M-N). Together our observations suggest a model wherein the peak of glycolysis in the tail bud/PSM imposes a high pHi to these cells thus favoring non-enzymatic β- catenin acetylation downstream of Wnt signaling. This could ultimately lead to activation of mesodermal transcriptional Wnt targets and specification of the paraxial mesoderm.

Our data is also consistent with posterior PSM and tail bud cells exhibiting Warburg-like metabolism with high aerobic glycolysis and an inverted pHe/pHi gradient as compared to more differentiated cells. Decreasing the pHi in Wnt-addicted tumor cells was recently shown to inhibit Wnt signaling, as reported here for differentiating tail bud cells (*26*). Thus our findings further emphasize the tight similarity between the developing tail bud cell and cancer cells that exhibit high Warburg metabolism resulting in high pHi and low pHe (*27*), supporting the notion that some tumor cells might reactivate a developmental metabolic program.

## ACKNOWLEDGEMENTS

We thank Norbert Perrimon and members of the Pourquié lab for critical reading of the manuscript and discussions. Y.H. acknowledges Grant-in-Aid for JSPS Fellows (29-456). Research in the Pourquié lab was funded by a grant from the National Institute of Health (5R01HD085121).

## AUTHOR CONTRIBUTIONS

M.O. designed, performed and analyzed the chicken embryo experiments and the *in vitro* acetylation study of purified beta-catenin with O.P.; Y. H. designed, performed and analyzed the human iPS experiments with O.P.; F.X. performed the quantitative analysis of pHi *in vivo*. O.P. wrote the manuscript and supervised the project. All authors discussed and agreed on the results and commented on the manuscript.

## METHODS

### Chicken embryo culture

Fertilized chicken eggs were obtained from commercial sources. Eggs were incubated at 38 °C in a humidified incubator, and embryos were staged according to Hamburger and Hamilton (HH) (*6*). We cultured chicken embryos mainly from stage 9HH at 37°C on a ring of whatman paper on agar plates as described in the EC culture protocol (*7*). Chemically-defined plates (3.5 mm petridish X25) were produced by combining an agarose solution (0.15 g agarose melted by heating in 25 ml ddH2O (MilliQ)), and 25ml 2XDPBS solution with 8.3mM, 0.83mM, and 0mM glucose. For chemically-defined medium, a 2X PBS solution (pH5.3, pH6.0, pH7.2, pH7.6) was prepared by combining 50 ml 10XDPBS (Sigma; D1283) and 450ml of ddH2O, adding (0g: pH5.3, 0.15g: pH6.0, 0.6g: pH7.2, 1.5g: pH7.6) sodium bicarbonate (Sigma, S5761) and 2.5 ml Penicillin-Streptomycin solution (Gibco (10,000 U/mL)). Embryos were prepared on a ring paper as for EC culture and incubated on the agarose plates in 2 ml of chemically-defined medium. For drug treatments, 2mM 2DG (2-Deoxy-D-glucose; Sigma) and 10mM sodium acetate (Sigma) were added to the plates.

### Time-lapse microscopy and axis elongation measurements

Stage 9HH chicken embryos were cultured ventral side up on a microscope stage using a custom built time-lapse station (*28*). We used a computer controlled, wide-field (10× objective) epifluorescent microscope (Leica DMR) workstation, equipped with a motorized stage and cooled digital camera (QImaging Retiga 1300i), to acquire 12-bit grayscale intensity images (492 × 652 pixels). For each embryo, several images corresponding to different focal planes and different fields were captured at each single time-point (frame). The acquisition rate used was 10 frames per hour (6 min between frames). To quantify axis elongation length, the last formed somite at the beginning of the time-lapse experiment was taken as a reference point, and the position of the Hensen’s Node with respect to this somite was tracked as a function of time using the manual tracking plug-in in Image J (*29*).

### Whole mount in-situ hybridization

Stage 9HH embryos were cultured at 38 °C in chemically-defined medium. After 10 h of incubation, embryos were fixed in 4% paraformaldehyde (PFA). Whole mount *in situ* hybridization was carried out as described (*30*). Probes for *AXIN2* (*31*), *CMESPO* (*Mesogenin1)*(*32*), *SOX2* and *SAX1* (*33*) have been described.

### Plasmid preparation and electroporation for chicken embryo

The ratiometric pH sensor pHluorin (*4*) was used to generate the expression vector pCAGG- pHluorin-IRES2-Td-Tomato. Full length pHluorin sequence was sub-cloned in pENTR-1A vector (Invitrogen), and inserted into a pCAGGS-IRES2-tdt-RFA destination vector using the Gateway system (Invitrogen). Chicken embryos ranging from stage 6HH to stage 7HH were prepared for EC culture. A DNA solution (1.0-5.0 μg/μl) was microinjected in the space between the vitelline membrane and the epiblast at the anterior primitive streak level which contains the precursors of the paraxial mesoderm. *In vitro* electroporations were carried out with five successive square pulses of 8V for 50ms, keeping 4mm distance between anode and cathode using Petri dish type electrodes (CUY701P2, Nepa Gene, Japan) and a CUY21 electroporator (Nepa Gene, Japan).

### Intra-cytoplasmic pH measurement for chicken embryo

In order to measure the intra-cytoplasmic pH (pHi) in PSM and tail bud cells, stage 4-5 HH chicken embryos were electroporated with the pCAGGS-pHluorin-IRES2-Td-Tomato construct which contains the ratiometric pH biosensor pHluorin (*4*) and cultured until stage 12 HH at 38 °C. Embryos strongly expressing the constructs were subsequently selected based on Td- Tomato expression in the PSM and tail bud. We first tested whether pHluorin can accurately report on pHi differences when electroporated in the chicken embryo. Embryos were electroporated at the anterior primitive streak level with the pHluorin construct to target tail bud and PSM cells as described previously (*34*) and they were reincubated until they reached stage 12 HH. Embryos were then incubated in different buffers (pH5.5, pH6.5, pH7.5) from the pH calibration buffer kit (molecular probe) with 10 μM of the protonophores Nigericin and Valinomycin at 37 °C for 30 minutes. Exposure to the protonophores allows the cells’ pHi to equilibrate to the buffer pH. Then embryos were mounted on MatTek glass-bottom dishes soaked in pH calibration buffer (pH5.5, pH6.5, pH7.5). Images were captured using a laser scanning confocal microscope (TCS SP5; Leica or LSM780; Zeiss) at 37 °C in humidified atmosphere. The protonophore treatment completely abolished the gradient of pHluorin 488/405 signal ratio and the average emission signal ratio under 488/405nm excitation decreased as pH increased (fig. S3), indicating that the pHluorin reporter electroporated *in vivo* can report on intra-cytoplasmic pH changes. However, because it is impossible to perform the different steps required for calibration in the same embryo, different electroporated embryos were used for each pH value and for experimental measurements. Therefore, the absolute value of the pH could not be defined (Fig. 1A). Thus, for all *in vivo* measurements, we only compared the 405 to 488 nm fluorescence ratios.

To measure the pHi *in vivo*, embryos were electroporated at stage 4-5HH with the pHluorin construct and cultured in EC culture until stage 9HH and then cultured for 10 hours with and without 2DG in EC cultures or in chemically-defined conditions before mounting. Next, embryos were mounted on MatTek glass-bottom slides on a thin albumin/agar gel in the same culture conditions. Images were captured as described above. After image capture, 3-channel z-stacks (.lif) of individual embryos were exported using Fiji into single channel single plane images (.png) for import into GoFigure2 software (www.gofigure2.org, (*35*)). Spherical segmentations of 9 µm radius were generated manually on the Td-Tomato channel image to encircle single cells on the raw images. Intensities of the 405nm-excited and 488nm-excited pHluorin signals were automatically calculated within the segmentations by GoFigure2. The measurements were exported for further analysis in custom made Matlab routines (Mathworks). To compare pH between different embryos, segmentations from individual embryos measured in the same day experiment were averaged to obtain a global ratio, followed by unpaired t-tests. To compare pH between different regions of individual embryos, segmentations were first normalized to the global ratio to minimize the contribution of embryo to embryo variation of overall signal intensity, followed by paired t-tests.

### Maintenance and differentiation of human iPS cells

The cell line NCRM1 (RUCDR, Rutgers University) was used for most human iPS cell experiments. For Fig. 3B, the previously described MSGN1-RepV reporter line was used (*36*). Human iPS cells were cultured on Matrigel (BD Biosciences)-coated dishes in mTeSR media (StemCells Technologies). Differentiation was performed as described in (*12*). Briefly, cells were plated at a density of 30,000-35,000 cells/cm2 in mTeSR supplemented with 10μM Y-27362 dihydrochloride (Rocki; Tocris Bioscience #1254). The next day (day 0 of differentiation), the medium was changed to DMEM/F12 GlutaMAX (gibco # 10565-018) supplemented with Insulin-Transferrin-Selenium (ITS, Gibco), 3 µM Chir99021 (Tocris #4423) and 0.5 µM LDN193189 (Stemgent #04-0074) (CL medium). At differentiation day 3, 20 ng/ml FGF (PeproTech #450-33) was added for an additional 3 days. After 6 days of differentiation, cells were changed to DMEM medium (gibco #11965-092) supplemented with 10 ng/ml HGF (Pepro Tech #315-23), 2 ng/ml IGF-1 (Pepro Tech #250-19), 20 ng/ml FGF and 0.5 µM LDN193189. After differentiation day8, cells were cultured with 15% KSR in DMEM supplemented with 2 ng/ml IGF-1 until day10.

### Culture medium preparation

To examine the effect of medium pH, cells were incubated in media buffered at different pH for 3 hours at differentiation day2. To adjust the medium pH, HCl and Sodium bicarbonate (Sigma #S5761) were used and the pH of the medium was measured using a pH meter (Mettler toledo) or pH indicator papers (pH6.0-8.1) (GE Healthcare Life Sciences #2629-990). Prior to experiments, the medium pH was equilibrated by incubation in 5% CO2 at 37°C for 24 h.

To examine the effect of glucose-free condition, cells were incubated in glucose (-) medium for 6h at differentiation day2 or for 24h at differentiation day1. 5mM 2DG (Sigma #D8375) was used in 5mM D-(+)-Glucose (Sigma # G7021) containing DMEM medium (#D9807-02) to inhibit glycolysis. Culture in CL medium with 2 or 10 mM Sodium acetate (Sigma #S2889) for 24 hour at differentiation day1 was used to increase the acetylation level.

### Lactate assay

To detect lactate production by the differentiated iPS cells, we used a lactate assay kit (Bio Vision #K607-100) according to the manufacturer’s instructions. Measurement of O.D. at 560nm was performed using GloMax-Multi Detection System (Promega). Values were normalized by total protein amount in each well.

### Quantitative RT-PCR

RNA was extracted using Trizol (Invitrogen #15596-018), followed by precipitation with chloroform (Sigma #288306) and Ethanol (Sigma #459836). RT-PCR was performed using 1ng total RNA using SuperScript? Reverse Transcriptase (Invitrogen #18080-051). Following primers were used for PCR. Actin (Forward; ccaaccgcgagaagatga) (Reverse; ccagaggcgtacagggatag), LDH-B (Forward; ttgtggtttccaacccagtggaca) (Reverse; aaaatccatccatggcagctgctg), Msgn1 (Forward; ctgggactggaaggacagg) (Reverse; acagctggacagggagaaga), Tbx6 (Forward; aagtaccaaccccgcataca) (Reverse; taggctgtcacggagatgaa), Axin2 (Forward; ggagtgcgttcatggtttct) (Reverse; tgcatgtgtcaatggtaggg).

### Measurement of intracellular pH

The BCECF pH sensitive dye (ThermoFisher Scientific #B1150) was used to analyze the intracellular pH in human iPS cells according to the manufacturer’s instructions. After washing the cells with HBSS (gibco #14025-092), 1μM AM ester solution (diluted with HBSS) was added to the culture and incubated for 20 minutes at 37°C. Then, cells were washed twice with fresh culture medium and incubated for 10 min at 37°C. After replacing the medium with HBSS, the 488/405 nm ratio was measured using confocal microscopy (LSM780, Zeiss) or GloMax-Multi Detection System (Promega). To determine the absolute pH value, we incubated different wells from the same experiment in the calibration buffer (3 different pH PBS adjusted by HCl or NaOH) with 10μM of the protonophores Nigericin and Valinomycin at 37°C for 15minutes. Then, the 488/405 nm ratios were measured using confocal microscope or Glo-Max multidetection System and used to generate a calibration curve from which wells from the same experiments could be compared.

### Immunocytochemistry

Cells were washed with PBS and fixed in 4% PFA /PBS for 20 min at room temperature. Fixed cells were washed with PBS, then blocked and permeabilized with 3% FBS and 0.1% Triton X- 100/PBS at room temperature for 30 min. Cells were then incubated with an anti-β-catenin antibody (1/500, BD #610153) and a chicken anti-GFP antibody (1/800, invitrogen #ab13970) diluted in PBS containing 3% Fetal Bovine Serum (FBS) and 0.1% Triton X-100/PBS at 4°C overnight, and next with secondary antibodies conjugated with Alexa 488 and Alexa 594 (1:500, Invitrogen #A11039, #A11037) for 30 min at room temperature. Images were captured using a laser scanning confocal microscope (LSM780, Zeiss)

### Quantification of nuclear localization

Nuclear localization of β-catenin was measured in cultures immunostained with the anti-beta-catenin antibody using the Fiji software. First, we splitted the channels and smoothened the Hoechst staining images using the “Gaussian Blur” filter. Then, we binarized the image, cut the border of adjacent cells using “watershed” and eliminated the β-catenin expression in outermost of nucleus area using the “erode” filter. We next manually drew the contours of the nucleus and used the “analyze particles” tool to measure the signal intensity of β-catenin in each nucleus.

### Western Blotting

Cells were collected with lysis buffer (50 mM Tris-HCl pH 8.0, 100 mM NaCl, 5 mM MgCl2, 0.5% Triton-X100, proteinase inhibitor cocktail (#78443S), 250 U/ml Benzonase) and incubated on ice for 30 min. After addition of SDS (final; 1%SDS), the samples were boiled and the protein concentrations were measured using a protein assay kit (BIO-RAD). The protein solution (10μg-20μg) was boiled in sample buffer and then ran on 10% or 12.5% SDS-PAGE. After the protein was transferred from the gel to polyvinylidene fluoride membrane (Millipore), the membrane was immersed in buffer containing 5% skim milk at room temperature for 1h. Membranes were then incubated with the primary antibody (diluted in 5% skim milk) overnight at 4°C. The next day, membranes were washed with PBT (PBS, 0.1% Tween 20) and incubated with Goat anti-Rabbit IgG secondary antibody, Horseradish peroxidase (HRP) conjugate (1/1000-1/10000; Invitrogen #31460) or Goat anti-Mouse IgG secondary antibody, and HRP conjugate (1/1000-1/10000; Invitrogen #31430) for 1h at room temperature. Immunoreactive bands were visualized with ECL Blotting Reagents (GE Healthcare #RPN2109) or SuperSignal™ West Pico PLUS Chemiluminescent Substrate (Thermo Scientific #34577) and detected using a Kodak X-OMAT 200A processor. After detecting the bands, membranes were washed with PBT and incubated in stripping buffer (2% SDS, 62.5mM Tris-HCl pH6.8, 0.7% 2- Mercaptoethanol) for 30min at 50°C to remove the antibody. Subsequent stainings were performed after removing the antibody. We used an Anti Acetyl-β-catenin (Lys49) antibody (1/1000; Cell Signaling Technology, #9534S), Anti-Active-β-catenin antibody (1/1000; Millipore #05-665), anti-β-catenin antibody (1/1000; BD #610153), anti-actin antibody (1/5000; Millipore #MAB1501), anti laminB1 antibody (1/5000; abcam #ab16048) and anti GAPDH antibody (1/1000; abcam #ab125247)

For chicken experiments, stage 9 HH chicken embryos were incubated at 37°C in different culture conditions (different pH or glucose concentrations) as described above. After 8 h of incubation, the posterior end of embryos was dissected and pooled (3 embryos for each sample). Samples were prepared, and Western blots were performed as for human iPS experiments.

### Immunoprecipitation

2-day chicken embryos were cultured on agar plates for 8h as described above and lysed using the lysis buffer of the Immuno-precipitation kit (abcam: ab206996). Total protein concentration was adjusted to 100μg per sample. Each sample was immuno-precipitated using 1μg anti-β- catenin antibody (BD #610153) or 1 μg Normal Mouse IgG (ab188776) overnight at 4°C according to the manufacturer’s instructions. After 3 washes using the kit’s washing buffer, samples were diluted into 2X sampling buffer, then Western blotting was performed as described above using Anti Acetyl-β-catenin (Lys49) antibody (1/1000; Cell Signaling Technology, #9534S), Anti-acetyl Lysine antibody (1/1000; cell signaling #9441), anti-β-catenin antibody (1/1000; BD #610153).

### In vitro acetylation of β-catenin proteins

Recombinant human β-catenin protein (20 μg; abcam ab63175) was incubated with 500μM or 10mM acetyl-CoA sodium salt (Sigma; A2056) in PBS solutions (final volume 20ul) at different pH conditions for 3 hours at 37°C. PBS solutions were prepared from 10XDPBS (Sigma; D1283) adding different amounts of sodium bicarbonate (Sigma, S5761). After incubation, 10μl of each sample were run on a 10% SDS-PAGE gel. Western blot was performed as described above using Anti-acetyl Lysine antibody (1/1000; cell signaling #9441), and anti-β-catenin antibody (1/1000; BD #610153).

### Nuclear fraction

Nuclear and Cytoplasmic Extractions were performed using NE-PER™ Nuclear and Cytoplasmic Extraction kit (Thermo Fisher Scientific #78833) in accordance with the manufacturer’s instructions. After extraction, proteins were analyzed by Western blotting as described above.

### Statistical analysis

Statistical analyses were performed with Prism 7.0 software (GraphPad). *P* values lower than 0.05 were considered to be significant. Statistical methods used in the analysis were described in figure legends.

---

**Figure S1:**
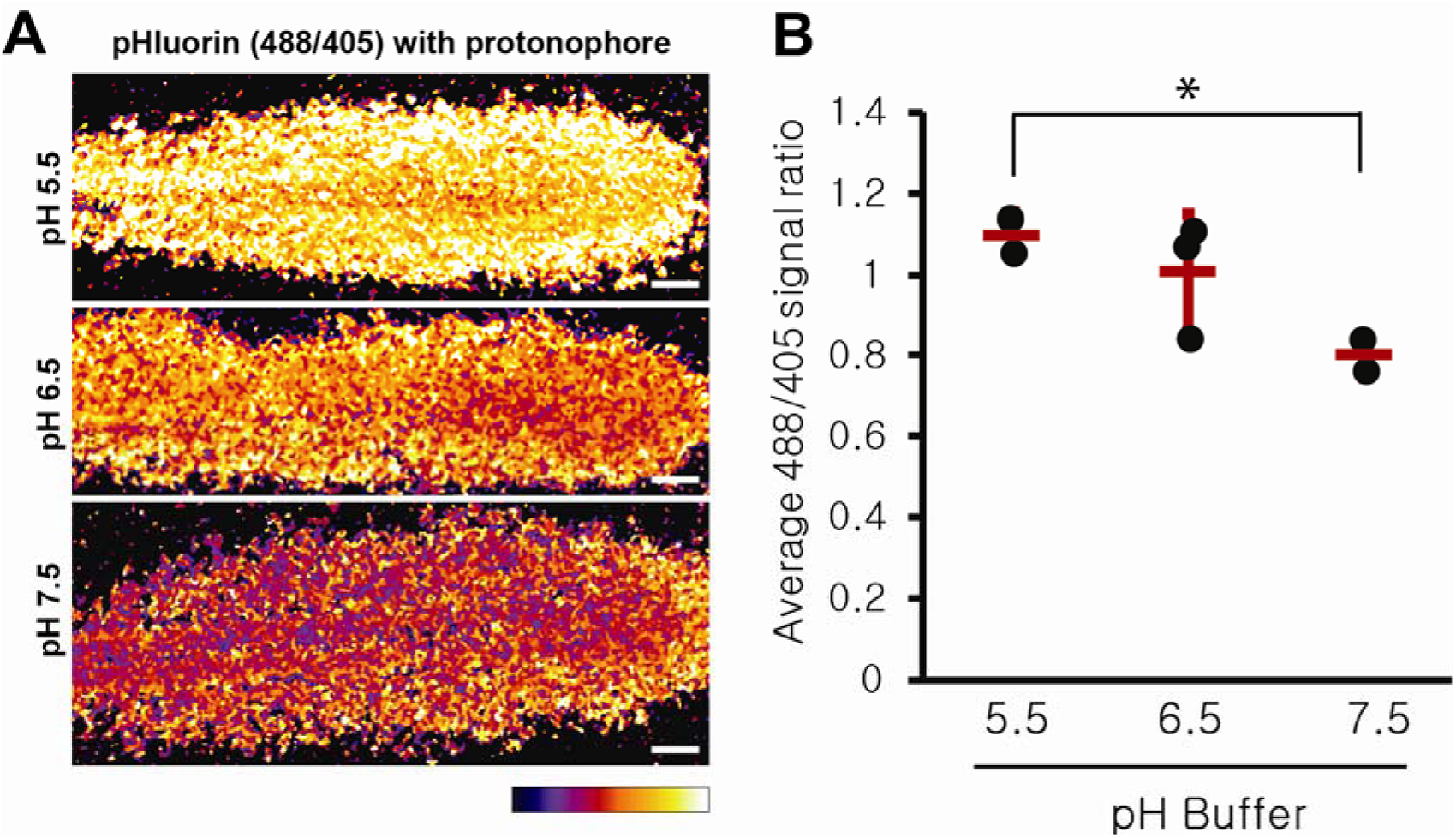
pHluorin can monitor intracellular pH in chicken embryos. (A) Ratiometric live expression of pHluorin (488/405 nm) detected in the posterior domain of electroporated embryos exposed to different pH buffers and Nigericin and Valinomycin. Fluorescence intensity is shown by pseudocolor image (Fire color) using image J. Yellow signal indicates lower pH. Ventral view, Anterior to the left. Scale bar :100 µm. (B) Each dot represents the average 488/405 nm signal ratio of ∼300 single cells segmented in one embryo (n=2 for pH 5.5, n=3 for pH 6.5, n=2 for pH 7.5.* p<0.05; t-test). Embryos were treated for 20 min in different pH buffers with the protonophores Nigericin and Valinomycin, before live imaging.

**Figure S2:**
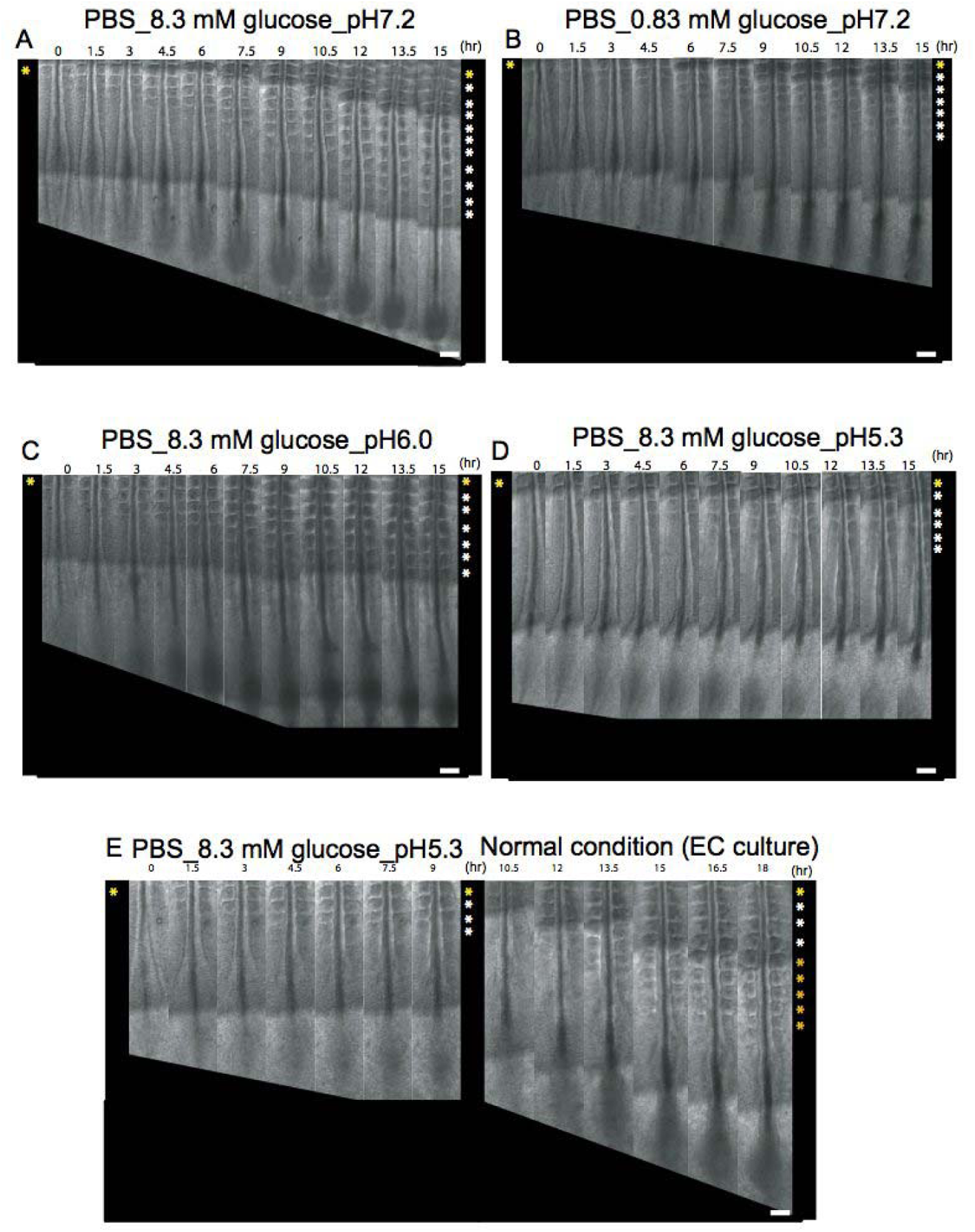
Axial elongation recovers when embryos are switched from acidic to normal pH medium. (A-B) Snapshots of 2-day chicken embryos cultured in minimal medium at pH 7.2 with 8.3 mM (n=6) glucose or 0.83 mM glucose (n=3). (C-D) Snapshots of 2-day chicken embryos cultured in minimal medium with 8.3 mM glucose in acidic conditions (pH6.0: n=6, pH5.3: n=6) (E) Snapshots of a 2-day chicken embryo first cultured in minimal medium with 8.3 mM glucose at pH5.3 showing the arrest of elongation after 9 hours and returned to control medium after 10.5 hours showing the rescue of elongation (n=6). Bright field micrographs of the posterior region of chicken embryos taken at 1.5 hour intervals. Somites formed at the last time point are indicated by asterisks on the right. Ventral views, anterior to the top. Scale bar, 100 μm.

**Figure S3:**
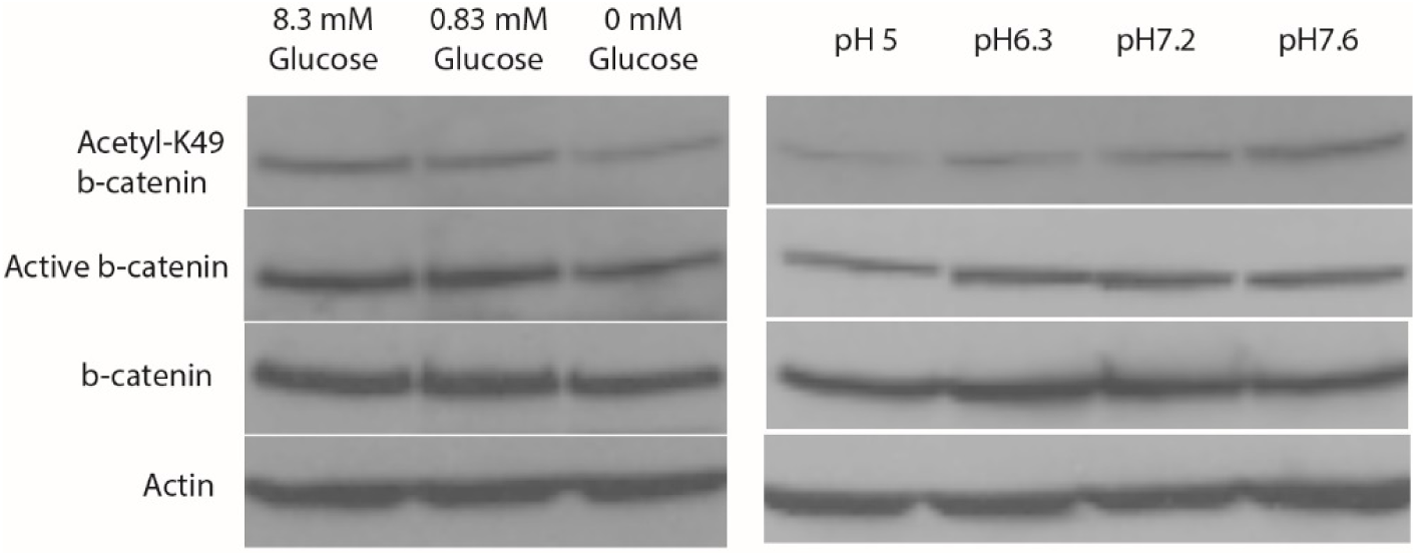
Acetyl-K49 β**-catenin expression level is affected by glycolysis activity and pHi**. Western blot analysis using anti-acetylated K49 β-catenin, anti-active β-catenin, anti-actin and anti-β-catenin antibodies. Extracts of 2-day chicken embryos cultured in minimal medium at pH7.2 with various glucose concentrations (n=4 per condition) or in minimal medium with 8.3 mM glucose at various pH (n=3 per condition).

